# Replacing continuous stimulation of one set of electrodes with successive stimulation of multiple sets of electrodes can improve the focality of transcranial temporal interference stimulation (tTIS), especially the focality of stimulation towards deep brain regions: A simulation study

**DOI:** 10.1101/2022.12.30.522357

**Authors:** Yixuan Li, Wei Zhang

## Abstract

Temporal interference (TI) stimulation has been proved to stimulate deep brain region while not activating the overlying cortex in the mouse brain. However, traditional TI, where two alternating currents with different frequencies are delivered by two electrode pairs, has poor focality in the human brain due to the high complexity of human brain structures, which limits the further application of TI. In traditional multipair TI, more than two alternating currents with different frequencies are injected by more than two electrode pairs at the same time. In this study, we proposed a novel multipair method which replaces continuous stimulation of one set of electrodes(eg: 1 montage for 30 minutes) with successive stimulation of multiple sets of electrodes(eg: 6 montages for 5 minutes each). By simulation using finite element method, We showed that this new method could improve the focality of TI in both neocortical regions and deep brain regions compared with traditional two-pair TI and the improvement was more significant in deep brain region. Ten realistic finite element models were used and five brain regions were tested(M1, DLPFC, ACC, NAc and hippocampus). It is expected that this new multipair TI can be used to improve the effectiveness of TI when researchers are translating TI from theoretical researches to clinical applications.

## Introduction

According to the global burden of disease study in 2016 and 2019, brain diseases, including both neurological disorders and mental disorders, has become the leading cause of disease burden all over the world since 1990 [1] [2] [3]. Drug therapy and psychotherapy are traditional treatment methods for brain diseases while physical therapy is a relatively new treatment and has great potentials [4]. Among all kinds of neuromodulation techniques, electrical stimulation has the longest history, the most tested safety experience and is most familiar to researchers [5]. There are two main kinds of electrical stimulation techniques, the first is deep brain stimulation(DBS), which needs a craniotomy and places electrodes in deep brain regions. DBS has been proved effective in Parkinson’s disease [6] [7], epilepsy [8] and depression [9]. However, being an invasive method, DBS is limited by the potential for surgical complications [10]. Another method is transcranial electrical stimulation (tES), including transcranial direct current stimulation (tDCS) and transcranial alternating current stimulation (tACS). tDCS can increase the excitability of neurons near the anode and decrease the excitability of neurons near the cathode [11] while tACS is believed to entrain the endogenous brain oscillations [12]. If you keep increasing the current intensity, tDCS and tACS can deliver a large amount of electrical field to deep brain regions but the inducing of larger amount of electrical field in the overlying cortex is inevitable due to physical laws [13].

Recently, a new method called transcranial temporal interference stimulation (tTIS or TI) was proposed to stimulate deep brain region while not activate overlying cortex [14]. This method use two electrical pairs to induce two alternating current of different high frequencies with a small difference (eg: 2000Hz and 2020Hz) to form a spindle wave of low frequency (eg: 20Hz) in deep brain regions [14]. TI has been proved in both mice [14] [15] [16] and humans [17] [18] [19] [20] [21]. However, TI has bad focality in humans because of the complicated structure in human brains [22] [23]. To improve the focality of TI, multipair TI which use more than two electrode pairs to deliver more than two alternating currents with different frequencies has been proposed [23].

In this study, we come up with a new stimulation scheme, using successive stimulation of multiple sets of electrodes to replace continuous stimulation of one set of electrodes. We prove this scheme is better than one pair TI by FEM simulation. We use the same FEM software package, developed by ourselves, as the our previous study [17]. Our result of 5 brain regions(M1, DLPFC, ACC, NAc and hippocampus) is a lot better than a previous study of traditional multipair TI [23], which suggests this new scheme maybe more advantageous in improving the focality of TI.

To sum up, TI is expected to become the first non-invasive deep stimulation in the world, but the poor focality of TI technology hinders its further development. Our paper proposes one possible solution. Previous researchers, for example, used only 1 montage configuration for 30 minutes of continuous stimulation. If we use 2 montages configurations for 15min each, or 3 montages configurations for 10min each, or 6 electrode configurations for 5min each, then it is possible to improve the focality. Because it is possible that the six montages have a complementary effect, that is, they all activate the target brain region while they activate different off-target brain regions.

## Methods

In this study, data of 10 subjects scanned by 3TGE MRI machine at the central campus of USTC in 2021 were adopted. The scanning sequences were 3D BRAVO and SAG 3d CUBE. Scanning instrument 3.0 Tesla MRI scanner with 8 channel head coil (Discovery MR750, GE, USA).

After obtaining MRI images, we used CAT12 module in SPM12 software package of MATLAB to register images, divide tissues, perform Boolean operations, and divide triangles on the surface. We use Tetgen library to construct tetrahedral mesh, obtain tetrahedral mesh with Delaunay property, and repair the four-point coplanar situation. So here we have a personalized tetrahedral grid of each subject’s brain. We use EEG 10-10 system, which contains 76 possible electrode positions.

Then, we use a GPU-accelerated software developed by ourselves to calculate the optimization problem of TI, which is, given the target brain region, to find the best montage to stimulate the target. We sort all montages in descending order according to ratio defined as formula 1, where ROI means region of interest, other means the brain except ROI:

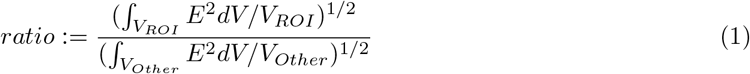

For each subject, we will get a ranking of montages in descending order of ratio. As stated in introduction, past researchers used only the top-ranked electrode combination to stimulate for a certain amount of time, say, 30 minutes. Below we show that replacing continuous stimulation of one set of electrodes with successive stimulation of multiple sets of electrodes can improve the focality of TI.

To save computation overhead, we use a heuristic algorithm to find complementary montages instead of exhaustive algorithm. We know the electric field intensity of each small tetrahedron of the first montage in the original list and, in Other area, classify the area with E greater than 0.2V/m as penalty and the area with E less than 0.2V/m as irrelevant. We re-definite ratio as formula 2:

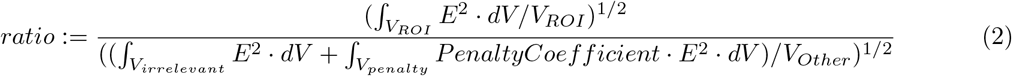

and use this new ratio to calculate a new ranking of montages. In this new ranking, because penalty coefficient is greater than 1, the top electrode combinations will stimulate the penalty region less.

The introduction of penalty coefficient is the most important step of our heuristic algorithm. In the exhaustive method, we just aimlessly make pairs of montages; In the heuristic algorithm, due to the introduction of the penalty coefficient, we will pair the first montage with the montage that is more likely to complement it.

After selecting a new montage, we add it to montages which are already chosen according to the formula 3, where n is the number of selected montages and *E*_*old*_ is the arithmetic average of the electric field intensity chosen montages. Adding in this way ensures that all montages contribute to E equally.

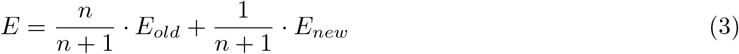

Our heuristic algorithm stops when the number of tetrahedrons which has a E larger than 0.2 V/m no longer falls or the number of montages reaches 10.

## Results

### Hot Graph

In Figure 1, we present the result of a single subject. The ROI is ACC. The MNI coordinate of the slice is chosen to near the center of the ROI. As we can see, the area beyond threshold (always chosen as 0.2 V/m in the field of tES simulation) drop a lot from pre to post. Beyond this qualitative result, in the next section we will show that the volume beyond threshold drop around 50% in the average result of 10 subjects.

**Figure 1.**
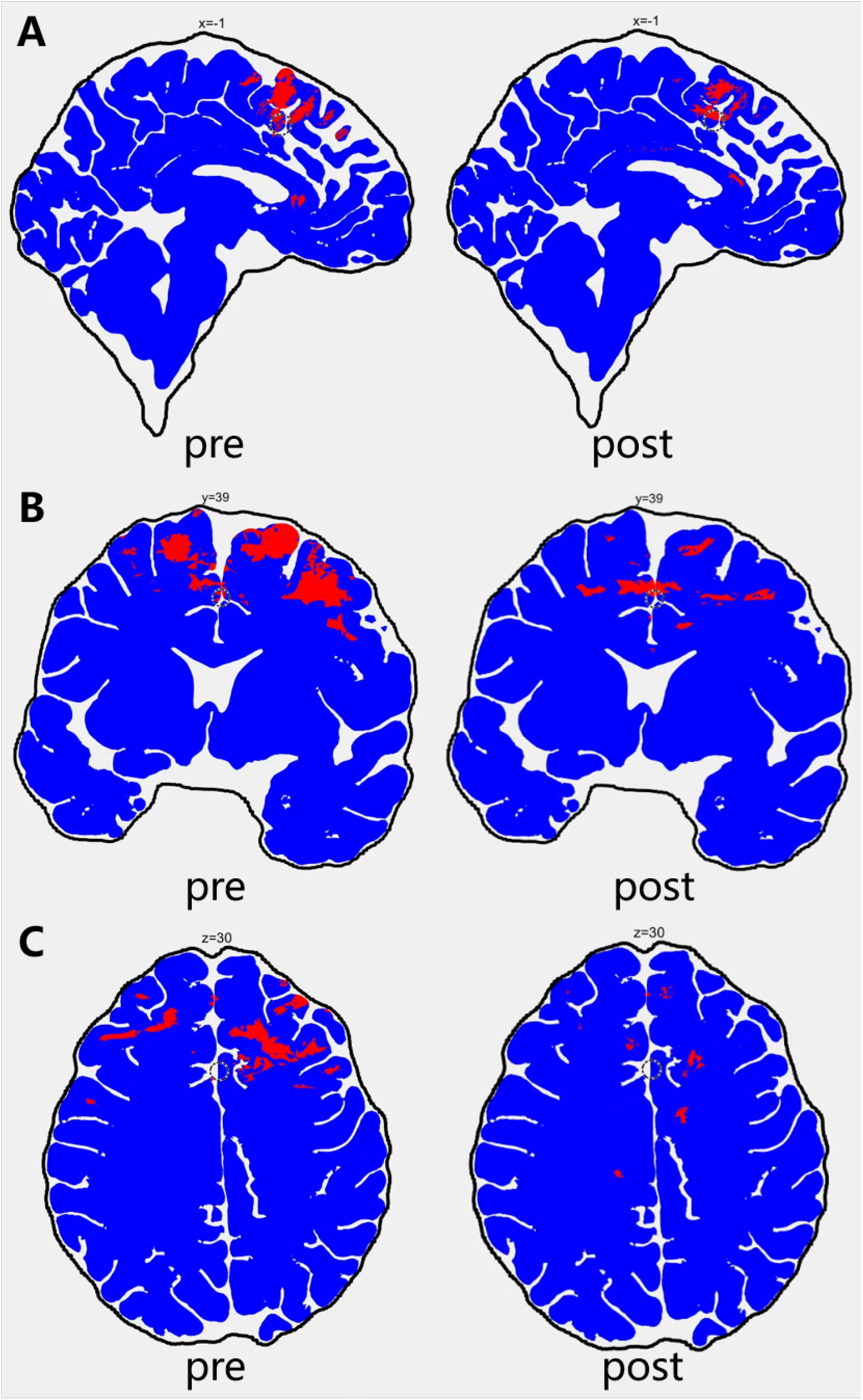
The circle indicates the ROI. A: sagittal plane, B: coronal plane, C: horizontal plane. Pre means the first montage of the original ranking. Post means 10 montages together. The blue area is *E <* 0.2*V/m* while the red area is *E >* 0.2*V/m*.

### Percentage

ROI means region of interest while other means the brain except ROI. We can see that this method is more useful in deep brain regions like ACC, NAc and hippocampus because the Other post/Other pre drops a lot while ROI post/ROI pre is nearly the same. To save computation time, we use the number of tetrahedrons as an approximation of volume. Because the volume of tetrahedrons is nearly the same, the approximation is acceptable. The MRI profile of these 10 subjects can be found in our GitHub page, see Acknowledgments.

How to interpret table 1? For example, in the result of the NAc, we sacrificed the 10% ratio in exchange for the number of small tetrahedrons of Other whose E beyond 0.2V/m decreased by 53%, and the number of small tetrahedrons of ROI whose E beyond 0.2V/m decreased by only 3%.

**Table 1.** Result of M1, DLPFC, ACC, NAc and hippocampus in 10 subjects. Pre means the first montage of the original ranking. Post means 10 montages together. Other post means the number of tetrahedrons beyond 0.2 V/m in the post situation. Other pre means the number of tetrahedrons beyond 0.2 V/m in the pre situation.

|  | Other_post/Other_pre | ROI_post/ROI_pre | ratio_post/ratio_pre |
| --- | --- | --- | --- |
| M1 | 31% | 84% | 70% |
| DLPFC | 45% | 80% | 89% |
| ACC | 57% | 96% | 92% |
| NAc | 47% | 97% | 90% |
| hippocampus | 44% | 89% | 92% |

Table 1 provides the simplest and the most intuitive data processing, demonstrates that replacing one set of electrodes with multiple sets can lead to very little change of the stimulation effect of ROI but make the stimulation effect of Other a lot worse.

However, the table cannot tell us the distribution of the number of small tetrahedrons in the brain with the intensity of electric field. In the next section we will show the histogram of the number of small tetrahedrons versus E, which will demonstrate why our method is useful.

### Histogram

In this section, we show the average of 25 subjects. All their MRI profiles can be found in our GitHub page too. The ROI is NAc.

Figure 2 shows that: compared with pre, the electric field in post showed the result that both sides decreased and the middle increased. Intuitively, the electric field in Other is more symmetrical. This confirmed our suggestion that the 10 groups of electrodes might have a complementary effect, that is, they all stimulated the target brain region, while they stimulated different off-target brain regions

**Figure 2.**
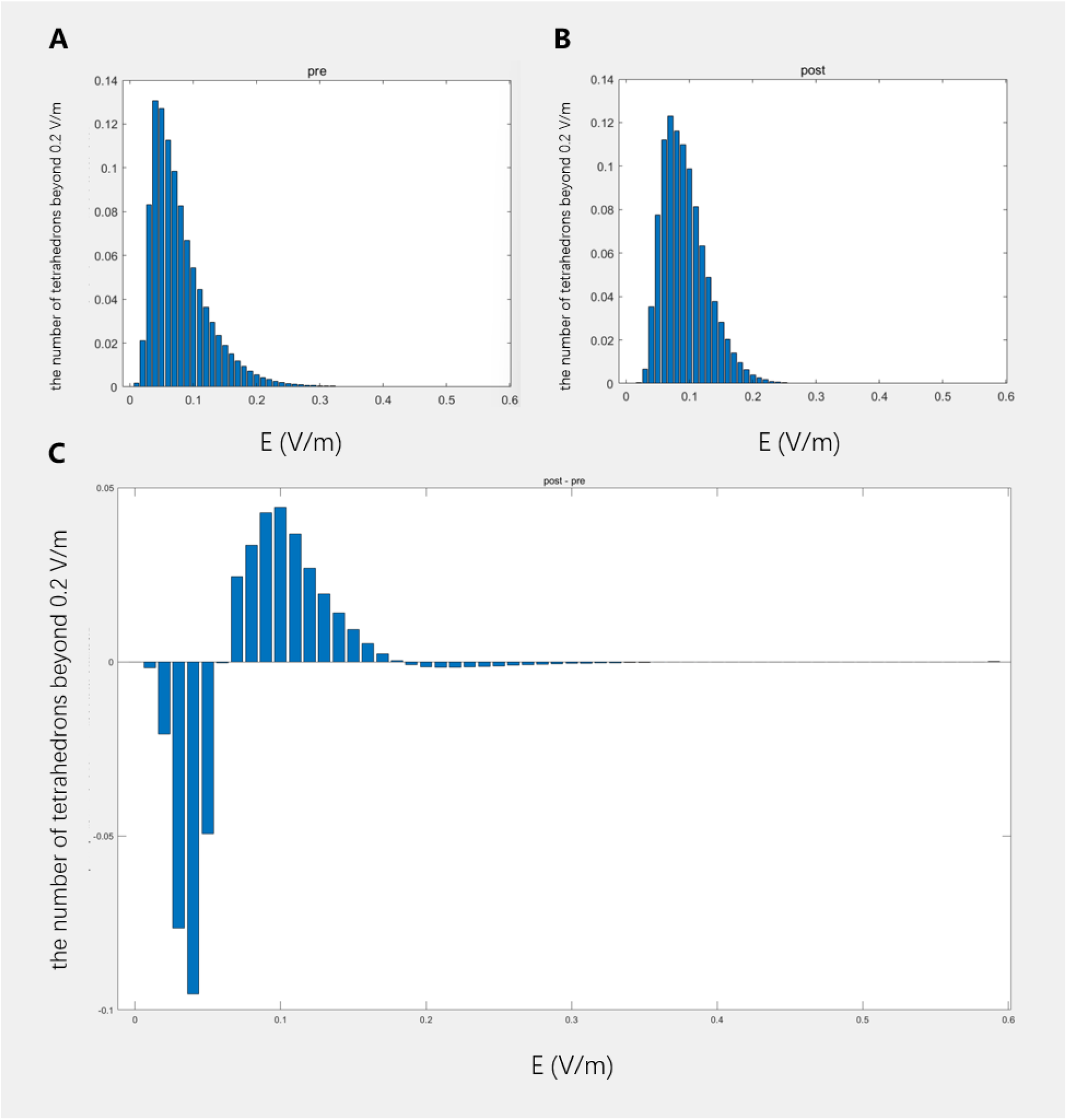
Result of NAc. A shows the distribution in pre, B shows the distribution in post, C shows post - pre

Figure 3 shows the distribution of the number of tetrahedrons beyond 0.2 V/m versus the distance to the center of the ROI. Compared with pre, in the post situation, the number of small tetrahedrons farther from ROI drops more. This indicates that the complementary effect of the 10 montages is better at the place far away from the ROI.

**Figure 3.**
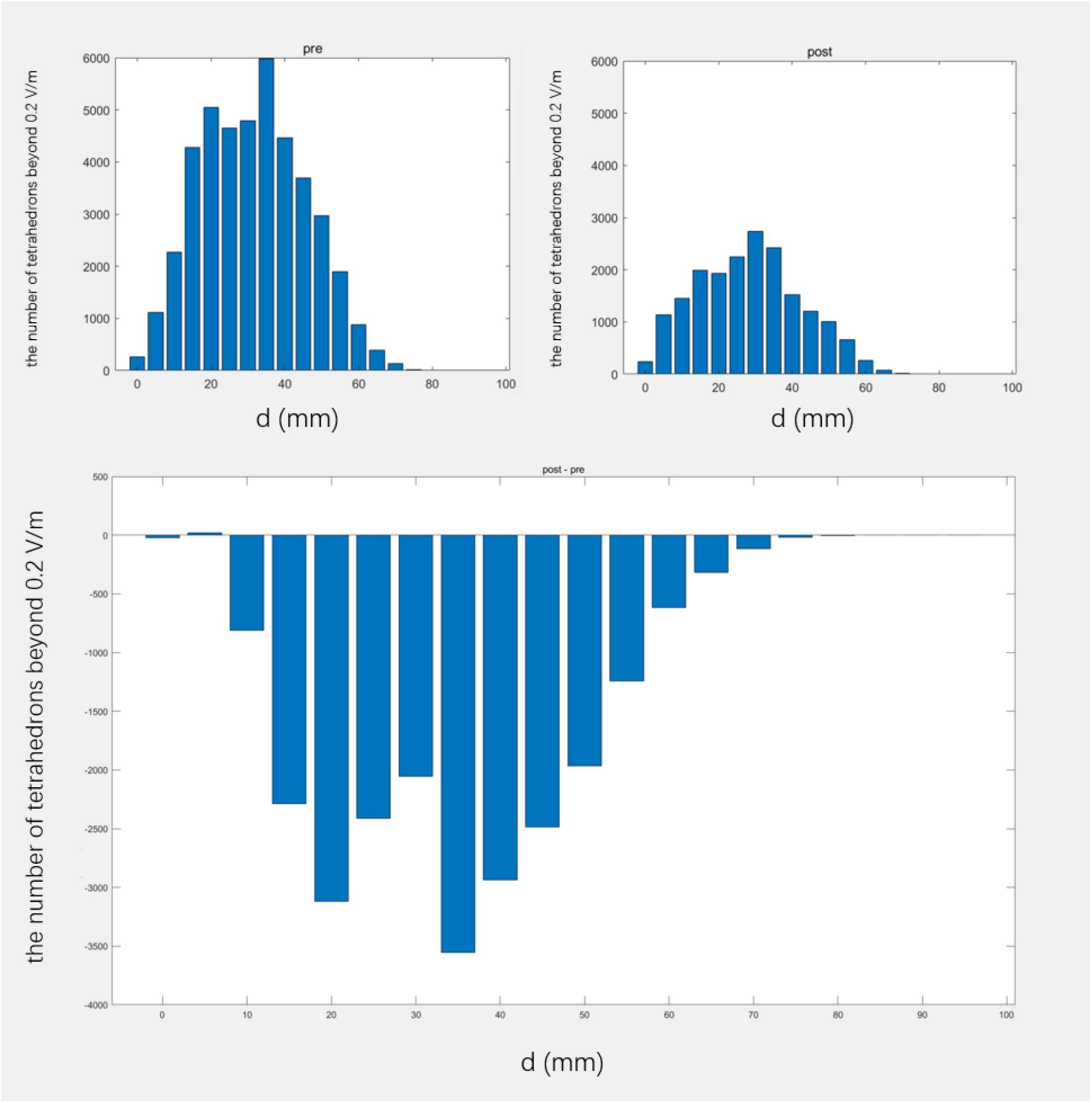
Result of NAc. A shows the distribution in pre, B shows the distribution in post, C shows post - pre. d means the distance to the center of the ROI.

Figure 4 shows the post - pre of another 3 brain regions (M1, DLPFC, ACC). They all show the same tendency as NAc. This means our method is robust under the change of ROI. Notice that Figure 4 is calculated use the average of 10 subjects (the same 10 subjects as previous section), not 25.

**Figure 4.**
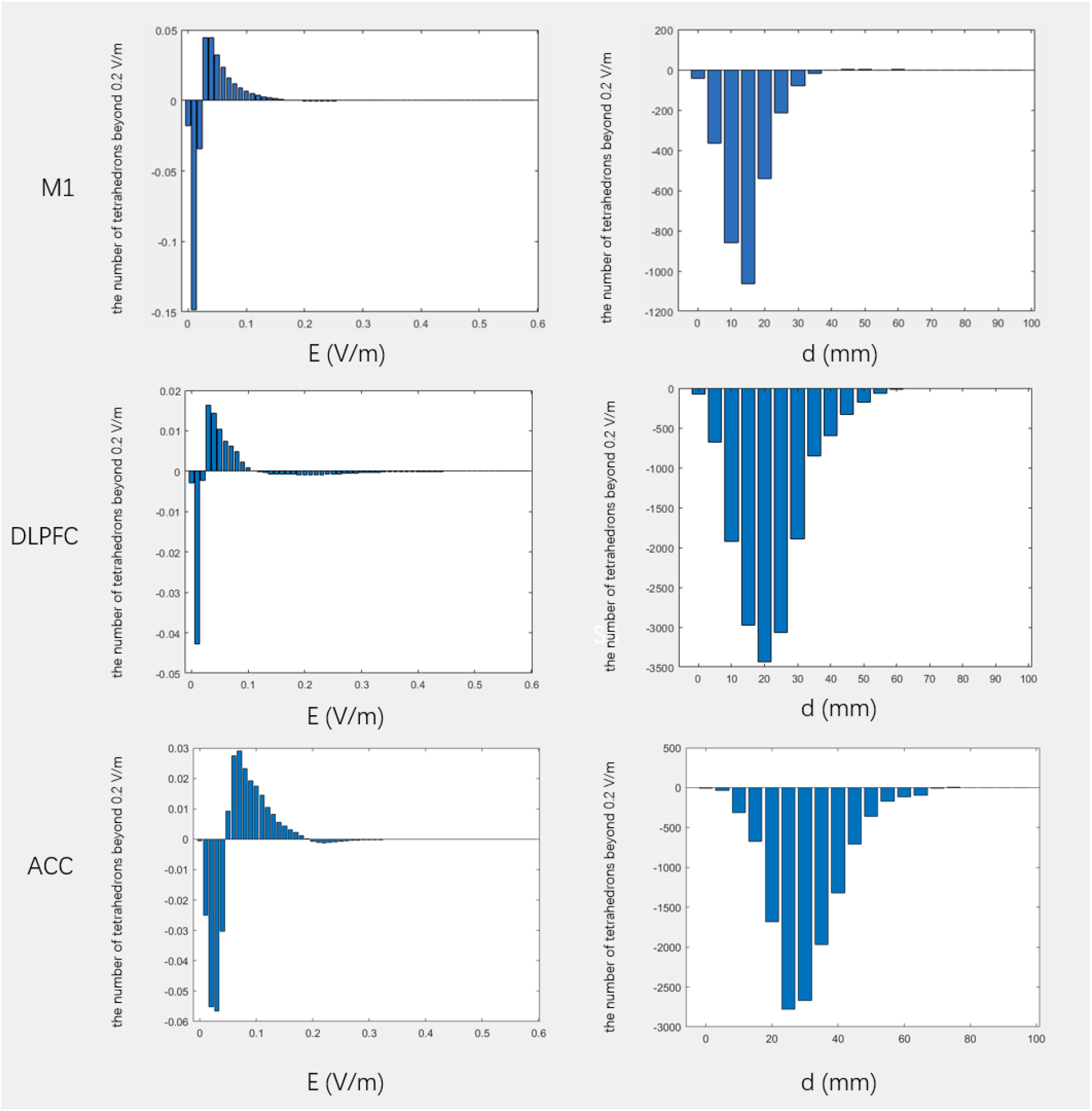
Result of M1, DLPFC, ACC.

In short, these histograms prove our conjecture: 10 montages do have a complementary effect in off-target brain regions while they all stimulate the target brain region well.

## Discussion

In this paper, we propose a new method, which replace continuous stimulation of one montage with successive stimulation of multiple montages and we show that it can improve the focality of TI, decreasing the volume beyond threshold to approximate 50%. Our method is very simple to implement in hardware, thus can be very useful when translate TI from theoretical researches to clinical applications.

The main drawback of our method is also the main drawback of TI: the max electric field intensity of TI (also of all tES methods) is only around 0.4V/m while the max electric field intensity of DBS can reach 75V/m. That is because the high conductivity of the scalp and skull and we can’t simply keep increasing the current intensity of tES, which will cause unbearable pain of subjects.

However, electric field intensity of 0.4V/m may still have some physiological effect, as shown in previous researches [17] [18] [19].

In short, our method is competitive stimulation scheme when researchers are translating TI from theoretical researches to clinical applications.

## Author Contribution

WZ first comed up with the idea. YX explored this idea, designed the simulation and wrote the paper.

## Acknowledgments

We thank everybody.

Our codes are available on GitHub, check them here.

